# PAG neurons encode a simplified action-selective signal during aggression

**DOI:** 10.1101/745067

**Authors:** Annegret L. Falkner, Dongyu Wei, Anjeli Song, Li W. Watsek, Irene Chen, James E. Feng, Dayu Lin

## Abstract

**Summary:** While the ventromedial hypothalamus, ventrolateral area (VMHvl) is now well established as a critical locus for the generation of conspecific aggression, its role is complex, with populations of neurons responding during the motivational, sensory, and action phases of aggression, and also during social interactions with the opposite sex. It has been previously unclear how the brain uses this complex multidimensional signal and generates a discrete action: the attack. Here we find that the largest posterior target of the VMHvl, the lateral periaqueductal gray (lPAG) encodes a simplified attack-selective signal during aggression. Single units in the lPAG exhibit greater selectivity for the attack action during aggression than VMHvl neurons and a subpopulation of neurons in the PAG exhibit short-latency, time-locked spiking relative to the activity of jaw muscles for biting during attack. In addition, channelrhodopsin assisted circuit mapping reveals a preferential projection from VMHvl glutamatergic cells to lPAG glutamatergic cells. Using projection-specific fiber photometry, we find that this excitatory projection conveys male-biased signals from the VMHvl to downstream glutamatergic PAG neurons that integrate ongoing male-related activity over several seconds, which suggests that action-selectivity is generated by a combination of both pre and postsynaptic filtering mechanisms.

---

The VMHvl has emerged as a clearing house for socially relevant information, responding during attack, but also exhibiting increased activity during sensory investigation of males and females and during the preparatory phase prior to attack(Falkner et al. 2016, 2014; Remedios et al. 2017; Lin et al. 2011). In addition, suppression of VMHvl activity decreases not only the frequency of attack, but also investigatory, sexual, and aggression-seeking behaviors(Yang et al. 2013; Lee et al. 2014; Falkner et al. 2016). How then do neurons downstream of the VMHvl interpret this complex code to drive aggression? Pharmacological manipulation of PAG circuits have been shown affect the efficacy of hypothalamic-mediated conspecific aggression, strongly indicating that the PAG’s actions are downstream of the hypothalamus (Zalcman and Siegel 2006; Gregg and Siegel 2003), though its precise role in this transformation has remained unclear. The emerging role of the PAG in the expression of other innate behaviors, such as stimulus-induced flight, appears to be that of a split-second action (Evans et al. 2018; Wang et al. 2019). We reasoned that a parallel circuit in the PAG might perform a similar function during conspecific attack.

We reversibly suppressed the activity of the PAG during freely moving interactions with males and females using the GABA_A_ agonist muscimol and tested whether inactivation altered conspecific social behaviors, including attack and investigation. We found that inactivation of the PAG produced action-specific deficits: we observed a decrease in time spent attacking a male intruder, but no effect in the time spent investigating either males or females (Fig 1A-B, Supplementary Movie 1). We observed a similar action-selectivity in the effects of muscimol on attack across alternative measures of behavior including mean duration of each behavior, latency to the first behavioral episode, number of behavioral episodes, and the total time spent engaging in each particular behavior (Supplementary Fig1). We did not observe a reduction of overall velocity during non-aggressive behaviors, suggesting that the effects of muscimol were not due to locomotor deficits (Supplementary Fig 1).

**Figure 1.**
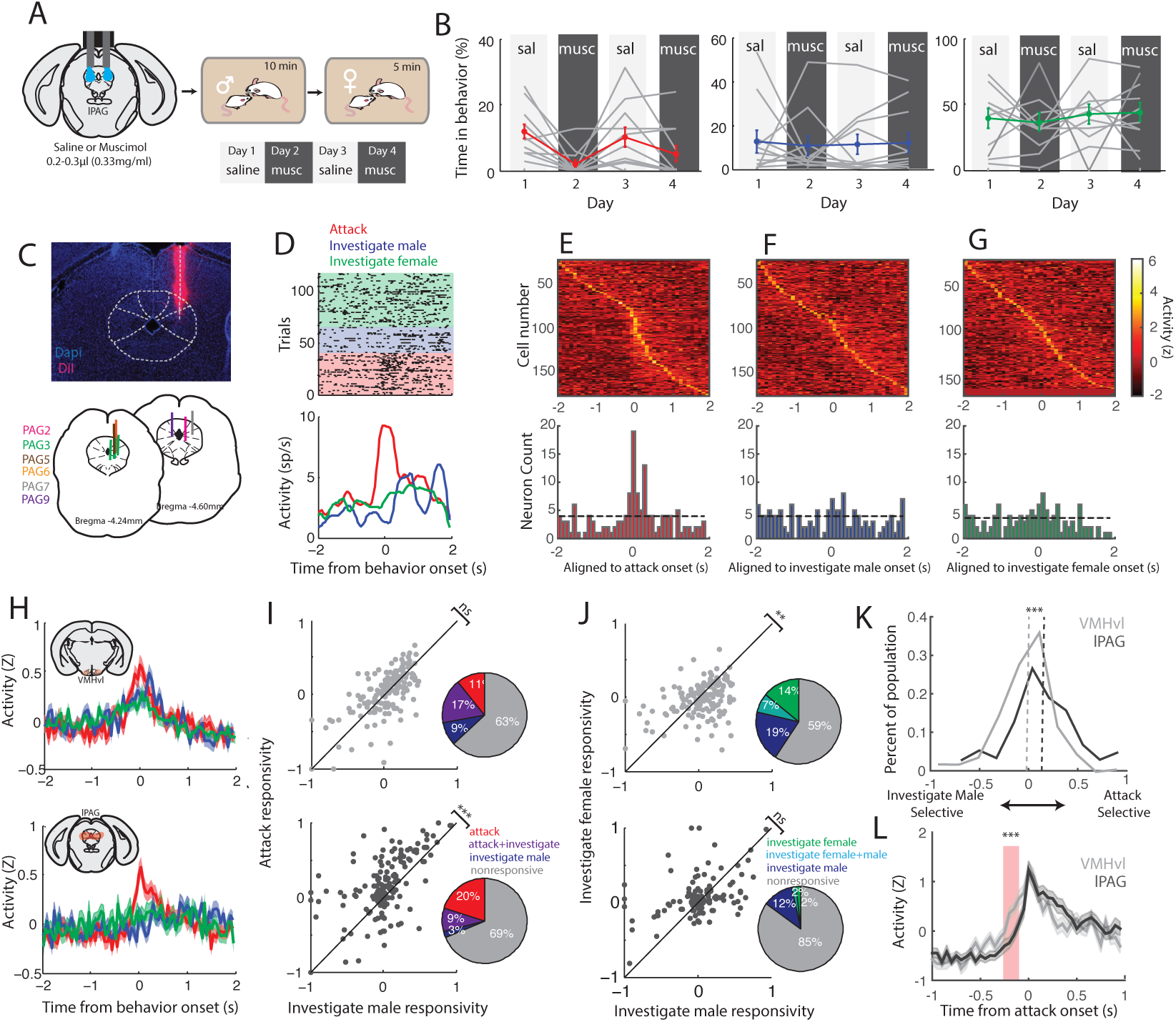
lPAG is more attack selective than the VMHvl. We recorded the behaviors during freely moving social interactions with males and females during reversible inactivation (A-B) and single unit recording (C-G). (B) Reversible inactivation of PAG using alternating injections of saline and muscimol induced selective deficits on attack and no change in investigatory behaviors (N = 11, *p=0.0134 for attack, left; p=0.985 for investigate male, middle; p=0.896 for investigate female, one-way repeated measures ANOVA). (C) Histology showing example placement of electrode bundle in the lPAG and electrode track locations for all recording animals (N=6). (D) Example raster plot (top) and PETH (bottom) for activity of an example lPAG units aligned to attack (red), investigation of a male (blue), and investigation of a female (green). (E-G) Normalized responses of population (top) of recorded neurons sorted by peak of response aligned to attack (E, N = 159), investigation of male (F, N = 158), and investigation of female (G, N = 151). Bottom histograms show number of units with response peak above 95% CI in each bin. Dotted black lines (E-G) represent chance levels for each behavior. (H) Normalized population response mean ±SEM of VMHvl (top) and lPAG (bottom) during onsets of key behaviors interactions with males: attack (red) and investigate male (blue), and with females: investigate female (green). (N = 166,156 for VMHvl male attack and investigate, N=212 for female investigate, N = 159 for lPAG male attack and investigate, N=151 for female investigate). Comparison of responsivity of individual VMHvl neurons (I-J top, light gray) and lPAG neurons (I-J bottom, dark gray). VMHvl population is nonselective between attack and investigate male (I, top, p=0.806, N = 157), and selective for investigation of male compared to female (J, top,p=0.005, N=147), while lPAG is selective for attack relative to investigate male (I, bottom, p=3.4×10^-7, N=152), and nonselective for investigation of males and females (J, bottom, p=0.415, N=152). Tests in I-J performed using Wilcoxon signed-rank test. Pie chart insets displaying percentages of individually significant neurons (Bonferroni-corrected t-test) in VMHvl and lPAG show an increasing number of purely attack selective neurons in the lPAG relative to VMHvl and a decrease of investigation selective neurons in the lPAG. (K) Selectivity of population to attack compared to selectivity to investigate male shows that attack-shifted peak for lPAG population (dark gray) relative to VMHvl (light gray). P=0.0001, Kolmogorev Smirnov test. (L) Attack responsive neurons in the VMHvl (light gray) have significantly increased activity prior to attack onset relative to attack responsive lPAG neurons (dark gray). N= 44 neurons in VMHvl N=46 neurons, lPAG, p = 0.0005, Bonferroni corrected unpaired ttest across all bins.

The specificity of this deficit during PAG inactivation indicates that the PAG’s role within the aggression circuit is uniquely action-selective relative to the more complex role of the VMHvl. To confirm this, we recorded populations of single neurons in the lPAG during free interactions with males and females and compared the response profiles to newly quantified measures of previously recorded neurons from the VMHvl (Falkner et al. 2014; Lin et al. 2011). We focused on the lPAG given the high amount of aggression-induced immediate early gene expression observed in our previous studies (Lin et al. 2011). Electrode tracks in all animals were confirmed to be located in the lPAG post hoc using DiI (Fig 1C). We recorded from populations of neurons during freely moving interactions with males (10 minutes) and females (10 minutes) and examined the firing rates of single units aligned to attack, investigation of males, and investigation of females (Fig 1D-G). Across the population (n=164 neurons, 6 animals), we observed that a subpopulation of neurons in the lPAG exhibited their peak firing aligned to the onset of attack while few neurons exhibited peak firing aligned to the onset of either investigation of male or female conspecifics (Fig 1E-G).

This selectivity for attack differs substantially from the response properties of the VMHvl. To directly compare these populations, we calculated the activity of each single unit in the VMHvl and lPAG to attack, investigate male, and investigate female. We observed that across the population, the VMHvl exhibits distinct activity peaks aligned to attack, investigation of males, and investigation of females (Fig 1H, top). In contrast, the population response at the PAG reveals an increase only during attack (Fig 1H, bottom). To compare the responses of single neurons in these populations, each neuron was assigned a responsivity score for each behavior (relative to its response during nonsocial behaviors). In addition, we determined the number of Bonferroni-corrected, significant neurons for each behavior in both VMHvl and lPAG populations (Fig 1I-J). We found that the population of lPAG neurons is significantly more attack-selective than investigate-selective, while the VMHvl population has equal responsive selectivity for both behaviors (Fig 1I). In addition, we found that lPAG exhibited an increased number of uniquely attack responsive neurons, and showed a decrease of both co-selective and investigate-selective single units relative to the VMHvl (Fig 1I, p=0.0031, chi-square test comparing numbers of attack responsive and investigate-responsive neurons between VMHvl and lPAG). Consistent with this, we also observed that the lPAG population shows little selectivity for either male or female investigation, and, relative to the VMHvl, has fewer neurons that respond to either or both of those behaviors (Fig 1J, p=0.0210, chi-square test comparing numbers of investigate-male-responsive and investigate-female-responsive neurons between VMHvl and lPAG). Overall, we found that the selectivity for attack (compared to other social behaviors) was increased in the lPAG compared to the VMHvl and that the activity of attack selective neurons in the VMHvl increased earlier than attack selective neurons in the lPAG (Fig 1K-L). Together these data are consistent with a role for the PAG in simplifying aggression-relevant signals from the VMHvl to drive attack.

To further explore the action selectivity of the lPAG during aggression, we examined the relationship between lPAG activity and an important effector for mouse aggression: the jaw. Using intramuscular injections of a retrogradely transported pseudorabies virus (PRV) (Smith et al. 2000), neurons in the PAG have been previously identified having polysynaptic projections to jaw muscles critical for aggressive behavior (Fay and Norgren 1997). We injected the superficial masseter muscle of the jaw (the critical muscle for jaw closure) with a GFP-labeled PRV 152 and found that the majority of retrogradely labeled neurons were located in the lPAG within the boundaries of the VMHvl excitatory terminal field (Supplementary Fig 2). We next tested whether these jaw projecting neurons were activated during aggressive encounters by examining the overlap between PRV-labeled GFP and the expression of an aggression-induced immediate early gene, c-Fos. We found that there were many c-Fos expressing neurons in the lPAG that were not GFP labeled, consistent with the idea that aggression may recruit many effector-specific pools of neurons. However, we also found that GFP-labeled neurons were far more likely than chance to also express attack-induced c-Fos (Fig. 2A-C), indicating that these jaw-projecting neurons in the lPAG likely play a role in aggression.

**Figure 2.**
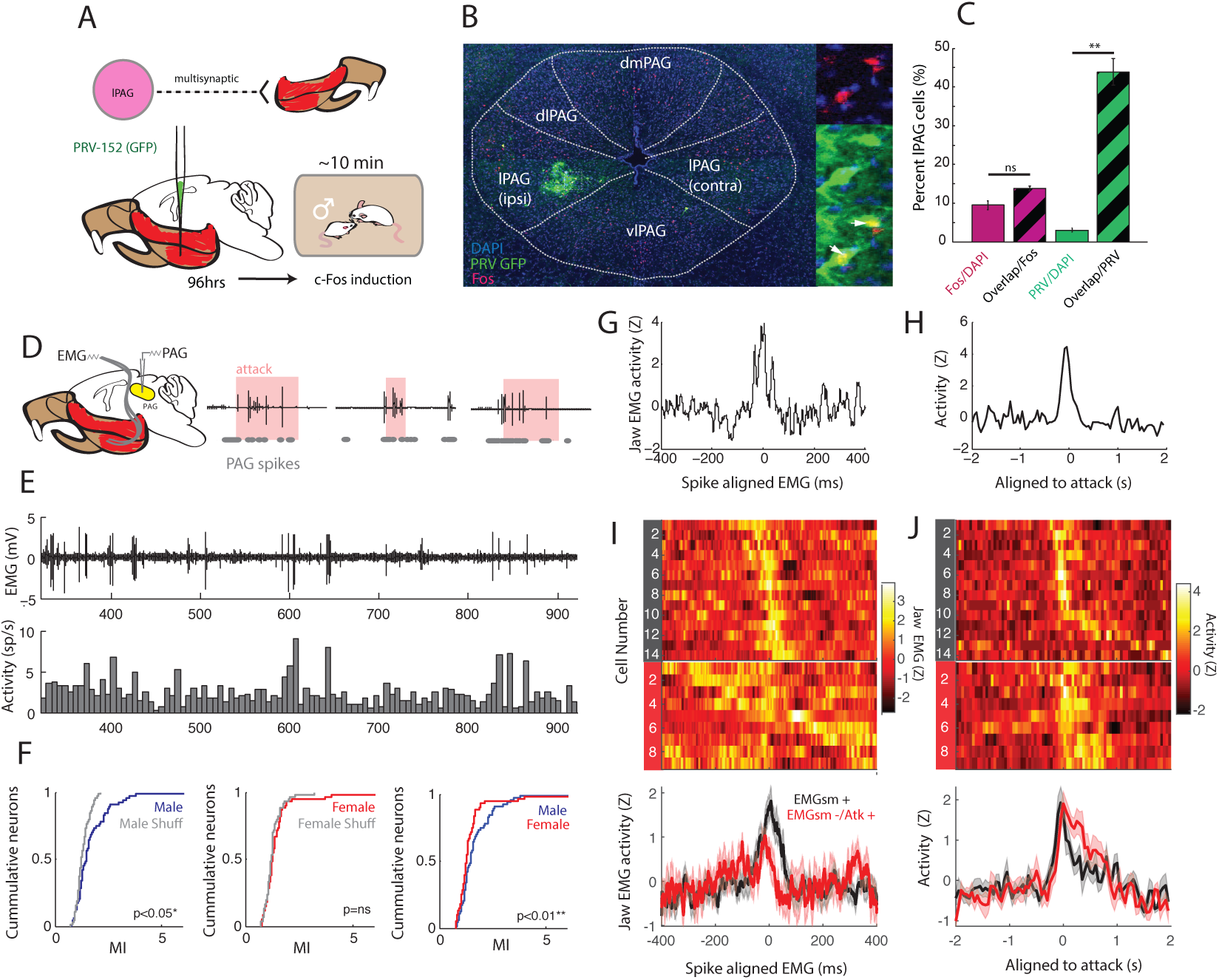
lPAG spiking has precise temporal alignment with jaw muscle activity during aggression. (A) Experimental design for injections of PRV-152 into the superficial masseter of the jaw, followed by 10min aggressive interaction for c-Fos induction after 96 hrs of viral incubation. (B) Example histology showing GFP labeling in the lPAG is co-localized with aggression-induced c-Fos. Panels show insets (white box) of c-Fos (mCherry, left top), PRV-GFP (left center), and merge (left bottom) with arrows indicating neurons with overlap. (C) PRV-GFP labeled cells show preferential overlap with aggression-induced c-Fos (p = 0.009, N = 3, paired t-test). (D) Example simultaneous recordings of jaw EMG and lPAG spiking during attack episodes. (E) Example EMG (top) and activity from simultaneously lPAG neuron (bottom) during interaction with a male (F) Mutual information (MI) of lPAG spiking and EMG activity comparing during interactions with males and females (N=64 neurons). MI of activity during male interaction compared to time shuffled control (p=0.0214, paired t-test), MI of activity during female interactions to time shuffled control (p=0.1551, paired t-test), MI of activity during male and female interactions (p=0.0053, paired t-test). Example STEMG (G) and attack-aligned PETH (H) with precise temporal alignment to EMG. (I-J) STEMG (I) and attack aligned activity (J) of neurons with significant STEMG (red, top trace), and significant attack responsive neurons that are do not have significant STEMG (black, bottom trace), show distinct dynamics.

We hypothesized that neurons in the lPAG may have a specific role in coordinating attack-relevant musculature. Since the time course of immediate early gene expression is too slow to resolve the temporal relationship between activity of PAG neurons and muscle activity, we developed a preparation to simultaneously record from neurons in the lPAG and EMG activity from the superficial masseter (EMG_SM_) during interactions with males while animals are attacking and performing other social interactions (Fig 2D-E, Supplementary Video 2, n=64 neurons in 4 animals). We used mutual information (MI) to quantify whether the EMG_SM_ signal provides useful information in predicting the activity during male or female interactions. Mutual information provides a model-free method for capturing the amount of joint information between two signals, and increased mutual information indicates that one signal can be used to predict the other (Srivastava et al. 2017). We calculated MI for each neuron and associated EMG_SM_ signal relative to a circularly permuted time shuffled control during interactions with males and females. We observed that across the population of recorded neurons, EMG_SM_ increased the MI during male interactions but not during female sessions relative to shuffle control and also that MI provided by the EMG_SM_ signal was significantly higher during male interactions compared to female interactions (Fig 2F), indicating that a subpopulation of neurons is modulated during attack-related jaw movement, but is not similarly activated by nonspecific jaw movements during female interactions.

We hypothesized that the activity of jaw-activated neurons might have a tight temporal relationship to the muscle activity if it is involved in directly activating the muscle. We examined the relationship between individual spikes and the EMG_SM_ signal. For each neuron, we generated a spike-triggered-EMG (STEMG) across an 800ms bin around each spike and used strict criteria to determine whether each STEMG demonstrated a significant relationship with individual spikes (Davidson et al. 2007). We found that a subpopulation of recorded neurons (21.8%, 14/64 neurons) showed a significantly increased STEMG within 60 ms of spikes during interactions with males (EMGsm+, Fig 2G and I, black). Most but not all of these EMGsm+ neurons (12/14) were activated during attack (Fig 2H and J, black). In addition, 9/64 neurons were identified as being significantly activated during attack but showed no increase in STEMG plot (EMGsm-/Atk+, Fig 2I-J, red). Those EMGsm-/Atk+ neurons often exhibited suppression following PAG spiking (Fig 2I, red) and showed activation that persisted through the attack responses (Fig 2J, red). These response patterns differ from those of the EMGsm+ cells that were increased only at the onset of attack (Fig. 2J, black). These data are consistent with the hypothesis that attack-related neurons in the lPAG may activate multiple attack related muscles, including jaw opening musculature. A smaller number of neurons (7.8%, 5/64 neurons) exhibited a time-locked relationship during female interactions, suggesting that some jaw responsive neurons may be recruited during other (nonaggressive) behaviors.

Though we did not explicitly test the activity of lPAG neurons during other nonsocial behaviors, we found that similar to the VMHvl, the population of recorded lPAG neurons exhibited decreased activity during spontaneous bouts of grooming and eating, indicating that non-aggressive jaw relevant actions do not similarly activate lPAG neurons (Supplementary Figure 3).

**Figure 3.**
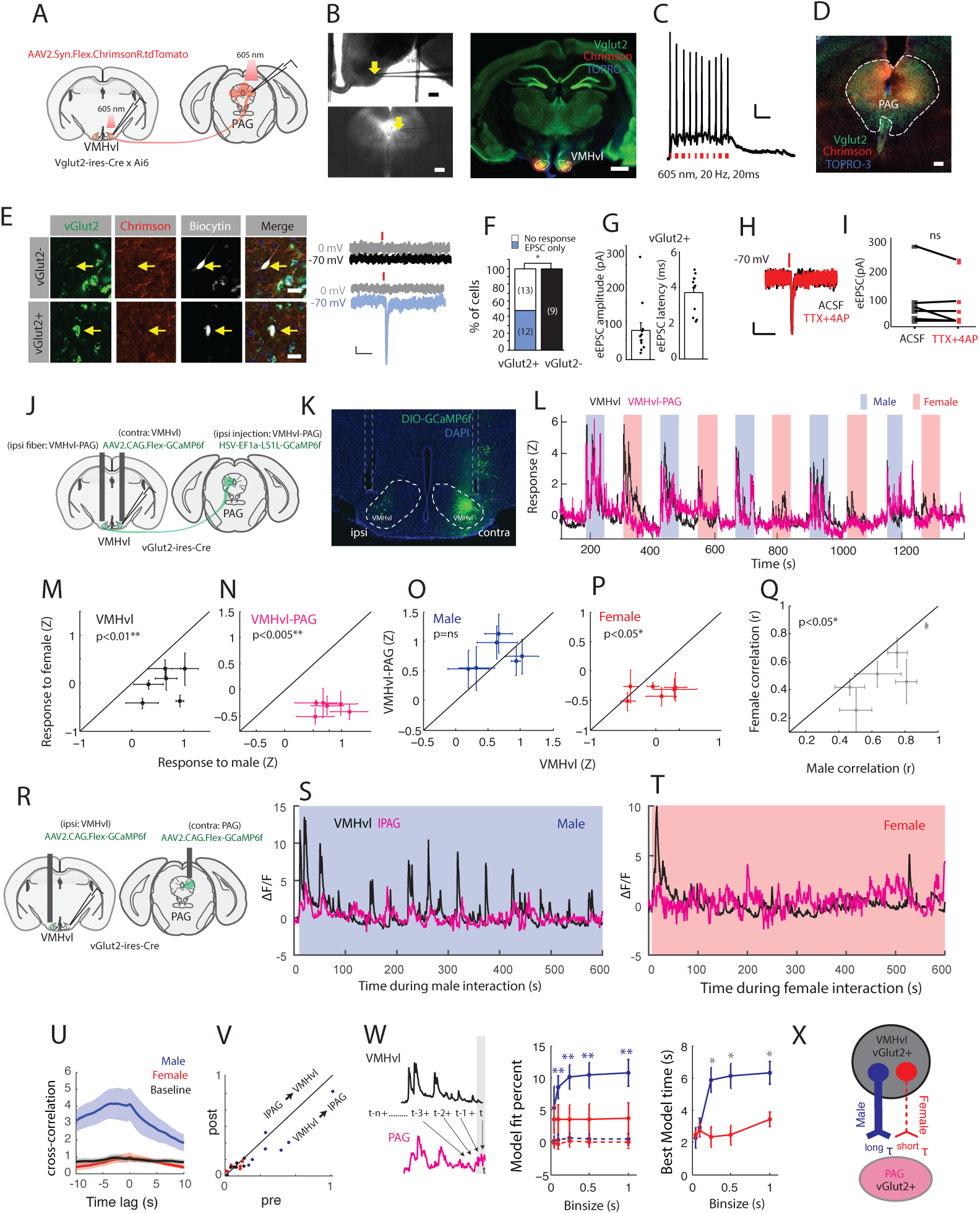
An excitatory VMHvl-lPAG projection filters aggression-relevant information using pre and post-synaptic mechanisms. (A) Viral strategy for targeting excitatory projections from VMHvl to lPAG using vGlut2-ires-cre x Ai6 mice. Slices were made of VMHvl and lPAG and whole cell recordings were performed. (B) Representative infrared differential interface contrast image (IR-DIC) from a recorded slice containing VMHvl (top, left) and lPAG (bottom, left). Yellow arrows indicate locations of recording pipette tips. Scale bar 500 µm. (right) Coronal section (right) showing expression of Chrimson-tdTomato (red) from a vGlut2 x Ai6 mouse. Scale bar 1mm. (C) Example trace showing current clamp recording of a VMHvl glutamatergic neuron expressing Chrimson-tdTomato. 605 nm light pulses (20 Hz, 20 ms for 500 ms, red ticks) reliably evoked time-locked spiking. Scale bars: 100 ms (horizontal) and 10 mV (vertical). (D) Histological image showing distribution of glutamatergic cells (green) and Chrimson-tdTomato expressing fibers from the VMHvl (red) in the PAG. Blue: Topro-3. Scale bar, 200 µm. (E) Enlarged views from (D) showing biocytin-filled vGlut2 negative (top row) and positive cells (bottom row) and their corresponding recording traces showing 1ms 605 nm light evoked EPSC (−70 mV) and IPSC (0 mV). Yellow arrows indicate the locations of the biocytin filled cells. Scale bar (left), 20 µm. Scale bars (right): 100 ms (horizontal) and 10 pA (vertical). (F) Stacked bar graphs showing the percentage of recorded PAG cells receiving EPSC during light stimulation. 12 out of 25 PAG vGlut2+ neurons received glutamatergic input from VMHvl, none showed light evoked IPSC; none of 9 vGlut2-neurons showed any light evoked responses. Fisher’s test. *p < 0.05. (G) Light-evoked EPSC amplitude (left) and latency (right) in PAG glutamatergic neurons (n=12). Error bars show means ± SEM. (H) Example traces showing 1ms 605 nm light evoked EPSC before (black) and after 1 µM TTX and 100 µM 4AP perfusion (red) in PAG glutamatergic neurons. Scale bars: 100 ms (horizontal) and 10 pA (vertical). (I) No change in light-evoked EPSC amplitude before and after 1 µM TTX and 100 µM 4AP perfusion in PAG glutamatergic neurons (n=6). Paired t-test. p > 0.05. (J) Experimental configuration of bilateral injection of Cre-dependent GCaMP6f into Vglut2-ires-Cre mice for simultaneous fiber photometry recordings of VMHvl and VMHvl-PAG projection neurons. Ipsilateral injection targets vGlut2+ VMHvl neurons and the contralateral injection targets vGlut2+ VMHvl-PAG projection neurons. (K) Example histology of GCaMP-labeled vGlut2+ VMHvl neurons (right) and Vglut2+ VMHvl-lPAG neurons (left) and placement of fiber tracks. (L) Example simultaneous recording of vGlut2+ VMHvl (black) and VMHvl-PAG (magenta) projection neurons during alternating interactions with males (blue) and females (red). (M-N) Population activity (mean +SEM for each animal, N=6 animals) of comparison between activity during male interaction and female interaction for VMHvl neurons (M, p=0.0062) and for VMHvl-PAG projection neurons (N, p=0.0002), shows that both populations exhibit increased activity to males. (O-P) Comparison of simultaneously recorded activity VMHvl and VMHvl-PAG neurons is not significantly different during male interactions (O, p=0.3676), but activity during female interaction is reduced in VMHvl-PAG neurons (P, p=0.044). (Q) Correlation of simultaneously recorded VMHvl and VMHvl-PAG neurons is higher in male interactions than female interactions (p=0.0238). All tests (M-Q) using paired t-test. (R) Experimental configuration of simultaneous recording of vGlut2+ populations in the VMHvl and lPAG. VMHvl and lPAG were recorded during interactions with males (S) and females (T). (U) Cross correlation of simultaneously recorded signals during male interactions (blue), female interactions (red) and no-interaction baseline (black). (V) Comparison of summed cross correlation in pre epoch (−10s to 0s) and post epoch (0s to 10s) for male, female, and no-interaction baseline shows significant asymmetry only during male interaction (N=7 animals, male p=0.0174*, female p=0.9333, baseline p=0.6148). (W) PAG activity was fit with a linear model using a variable amount of preceding VMHvl signal as the regressors. Fit percent (middle) and time (right) associated with best fit models of cross-validated data using a time-varying input from VMHvl. Dotted lines represent data from time shuffled controls. (X) Conceptual model of pathway selectivity of male-responsive information.

Our data demonstrates that neurons in the lPAG exhibit a greater degree of selectivity for aggressive action (i.e. attack) than neurons in the VMHvl. One simple mechanism by which this circuit could perform the transformation is if the projection from the VMHvl to PAG were a labeled line for aggression-specific information. To test this directly, first we explored whether excitatory projection neurons from the VMHvl form functional connections with neurons in the lPAG. We targeted excitatory projection neurons by injecting a red-shifted cre-dependent excitatory opsin (AAV2.Syn.Flex.ChrimsonR.tdTomato) into the VMHvl of vGlut2-ires-cre mice crossed with an Ai6 reporter. This strategy allowed us to record postsynaptic responses from vGlut2+ and vGlut2-neurons in the lPAG while optically manipulating the glutamatergic VMHvl-PAG projection (Fig. 3A-B). We first confirmed that brief light pulses at the VMHvl were sufficient to reliably evoke action potentials (Fig. 3C). Then, we made coronal slices of the lPAG and performed voltage clamp recording from putative vGlut2+ and vGlut2-neurons (identified by the presence and absence of GFP label) and tested whether a brief red light pulse activating the VMHvl-PAG terminals was sufficient to evoke postsynaptic responses (Fig. 3D-E). We found that approximately 50% (12/25) of identified vGlut2+ neurons exhibited short-latency excitatory post-synaptic currents (EPSCs) upon light delivery, while no single vGlut2-neurons (n =9) exhibited this excitatory response (Fig. 3F-G). Bath application of tetrodotoxin (TTX) and 4-aminopyridine (4-AP) did not change the magnitude of light-evoked EPSCs, supporting the monosynaptic nature of the connection (Fig. 3H-I). None of the recorded cells showed inhibitory postsynaptic currents (IPSCs) upon light delivery.

Next, to explicitly test what information is being conveyed by this excitatory hypothalamic to midbrain projection, we targeted excitatory VMHvl-PAG projecting neurons, by injecting the ipsilateral side of a vglut2-ires-cre male mice with a retrogradely transported, cre-dependent, calcium indicator (HSV-Ef1a-LS1L-GCaMP6f) into the lPAG. On the contralateral side of the brain to the injection site, we also injected a cre-dependent calcium indicator (AAV1.CAG.Flex.GCaMP6f.WPRE.SV40) in the VMHvl in order to compare the activity from the total VMHvl vGlut2+ population, not specified by projection. We positioned fibers over both the ipsi and contralateral VMHvl and used fiber photometry to simultaneously record respectively from the VMHvl-PAG projection vGlut2+ neurons and the VMHvl general vGlut2+ population during interactions with males and females (Fig 3J-K). We alternated between brief presentations of a male mouse stimulus (~1 min) with a female mouse stimulus (~1 min) with a 1 min break between presentations (Fig 3L) and compared the normalized activity between the VMHvl-PAG and VMHvl neuron populations across the entire set of interactions.

We found that both vGlut2+ VMHvl and VMHvl-PAG populations showed a strong bias towards male compared to female mean activity (Fig 3M-N). However, in a direct comparison of activity levels between VMHvl and VMHvl-PAG populations, we found that activity during male interaction was not significantly different between these simultaneously recorded populations (Fig 3O-P), while during female interactions, VMHvl-PAG activation decreased relative to the overall VMHvl population response. In addition, to track how well the two signals match, we computed the correlation (Pearson r) between the simultaneously recorded population activity of VMHvl-PAG and VMHvl neurons and found that correlation coefficients during male interactions were significantly increased during male interactions relative to female interactions across animals (Fig 3Q), indicating that the VMHvl-lPAG projection conveys a more faithful copy of population activity during male interactions than female interactions.

While we found that the VMHvl-PAG projection conveys male-biased information, we did not observe that this pathway conveys action-selective information during aggression (Supplementary Fig 4). To quantify this, we compared the PETHs of activity in the VMHvl vGlut2+ population and the VMHvl-PAG vGlut2+ projection during attack and investigation of males. We found that, similar to the single unit recording data, both VMHvl and VMHvl-PAG neurons also showed clear activation peaks aligned to both attack and investigation of males that were very strongly correlated at the level of the individual behavior episode (Supplementary Fig 4A-C).

Together these data demonstrate that an excitatory projection from the VMHvl to the lPAG conveys male-biased information to excitatory downstream popusltions, behaving as a labeled line for aggression-relevant information. While this labeled line projection effectively filters female-evoked signals, it does not filter non-attack signals during male interactions such as investigation. This suggests that further mechanisms are needed to transform VMHvl activity to an aggression-specific signal. One possibility is that the PAG is sensitive to slow temporal features in its inputs prior to activation during attack, including investigation-evoked increases that precede attack. To specifically model the interactions between the VMHvl and the PAG, we performed simultaneous recordings of glutamatergic population (vGlut2+) in both areas during free interactions with males and females (Fig. 3R-T). Qualitatively, we observed that the during male interactions, PAG activity peaks often followed VMHvl activity peaks, while during female interactions, signal dynamics were more independent (Fig 3R-T, Supplementary Fig 4D-F). To quantify these interactions, we computed the cross correlation of the simultaneously recorded signals separately for male interactions, female interactions, and a baseline no-interaction epoch. We found that the cross correlation during male interactions was strongly and asymmetrically skewed across a multi-second timescale, indicating that increases in VMHvl signals “lead” PAG signals during interactions with males, but not females (Fig. 3U-V). The integral of the cross correlation during the “pre” epoch (VMHvl leads PAG) relative to the “post” epoch (PAG leads VMHvl) significantly increased during male interactions across the population of recorded animals (n=7, p=0.0174, paired t-test) and not significantly different during either the interactions with females or no-interaction baseline period (p=0.9333, p=0.6148, paired t-test).

The slow temporal dynamics of the cross correlation during male interactions suggests that PAG activity is influenced by VMHvl activity stretching backwards in time for several seconds. To specifically quantify this temporal relationship between the VMHvl and lPAG, we modeled the ongoing PAG activity using a time-varying linear regressive model. We iteratively fit PAG activity with the activity of the VMHvl (VMHvl → lPAG) with a family of models with a variable number of regressors representing increasing numbers of previous time bins, used Akaike Information Criteria (AIC) to select the best model within each family, and cross validated this model on a separate data set for each recorded animal. We performed this model selection separately for male and female interactions and across a variety of bin sizes, ranging from 50 ms to 1 s. We found that cross validated fits for the best fit model were significantly better during male but not for female interactions relative to a time shuffled control. Importantly, this effect was consistent across a range of bin sizes (Fig. 3W), indicating that this variable made little difference in the model fit. In addition, we extracted the model order (number of time bins) for the best fit model for each animal for male and female interactions for each bin size. We found that the elapsed time associated with these best fit models was significantly longer for male interactions than female interactions (Fig. 3W, right), indicating that during male interactions, PAG signals are influenced by VMHvl signals stretching farther back in time.

On a conceptual level, this suggests that activity in the VMHvl increases for several seconds prior to activation of the PAG, an effect consistent the action specificity of the PAG. As a control, we also fit the reverse circuit model, (lPAG→ VMHvl) and found that both the fits, and model order are significantly decreased relative with the forward circuit model during male, but not female interactions (Supplementary Figure 5). Together, these results suggest a model by which during interactions with males, but not during interactions with females, the PAG receives VMHvl inputs conveying information about the sensory properties of the stimulus obtained during investigation, and this information is integrated over many seconds in order to drive attack behavior (Fig. 3X).

These findings provide evidence that the role of the PAG in the aggression circuit is to transform the complex sensory-motor and motivational signals of the hypothalamus into aggressive action and to coordinate the activation of effector-specific musculature, including the jaw. These results add to a growing literature that position the PAG as a critical initiator of innate behaviors that are determined by combinations of noisy sensory and state-dependent inputs (Koutsikou, Apps, and Lumb 2017). Since PAG neurons are active later and more acutely during attack behavior than its hypothalamic inputs from the VMHvl, it may serve as a behavioral initiation threshold, as has been suggested for other innate behaviors. We do not mean to suggest that the lPAG is “for” aggression: the PAG (and in particular hypothalamic to PAG pathways) have been implicated in many innate sensory-driven behaviors including but not limited to threat responsivity (Wang, Chen, and Lin 2015; Evans et al. 2018), prey capture(Li et al. 2018), itch(Gao et al. 2018), oro-motor coordination(Stanek et al. 2014), and social avoidance following defeat (Franklin et al. 2017). Additionally, the PAG has a well-documented role across species in the generation of vocalization, a behavior that also requires the integrations of social-sensory signals and the coordination of facial and laryngeal musculature (Kittelberger, Land, and Bass 2006; Holstege 2014). Here we add to this growing literature by elucidating the neural coding during conspecific attack and further hypothesize that the PAG is capable of orchestrating many complex innate behaviors by coordinating output to relevant muscles through sex-selective processing of slow temporal features in its inputs.

## Methods

### Animals

Experimental mice were sexually experienced, wild-type male C57BL/6N (12–24 weeks, Charles River), wild-type male Swiss Webster (12–24 weeks, Taconic), *vGlut2*-ires-Cre mice. Naïve vGlut2 x Ai6 mice were used for slice physiology and tracing experiments. Intruders were either group housed, sexually inexperienced BALB/c males or C57BL/6 females (both 10–30 weeks). Mice were maintained on a reversed 12-h light/dark cycle (dark cycle starts at noon) and given food and water *ad libitum*. All procedures were approved by the IACUC of NYULMC in compliance with the NIH guidelines for the care and use of laboratory animals.

### Behavior analysis and tracking

All freely moving behaviors were recorded using top and side GigE cameras using StreamPix (Norpix) and all videos were manually annotated for pre-identified behaviors and tracked for positional and velocity information using custom Matlab Software (https://github.com/pdollar/toolbox). Behaviors were manually classified as previously described (Falkner et al. 2014); individual behaviors included attack, investigation of males, investigation of females, mounting, eating, and grooming. All interactions were “resident intruder assay” (5-10 minutes of free interaction with male or female intruder), or alternating interactions with males and females (1 min each) separated by 1 min (Fig 3).

### Pharmacological Inactivation

For all pharmacological inactivations, double cannulae were implanted 0.5 mm above lPAG (coordinates: −4.24mm,A-P, +/−0.5mm M-L, -1.85mm D-V) from and 0.2–0.3 μl of either saline or of 0.33 mg/ml muscimol (Sigma) in saline were injected into the PAG bilaterally through the implanted double cannulae on alternating days. 8/11 animals were injected with fluorescent-conjugated muscimol (0.5 mg/ml) prior to perfusion to confirm injection coordinates and injection volume spread.

### Extracellular recording of freely moving mice

Methods for physiological recording in freely moving animals were described previously. Custom-built 16-channel (or 14 channel with EMG) tungsten electrode bundles or groups of tetrodes were attached to a moveable microdrive and implanted over the lPAG. After allowing 2 weeks for recovery, we connected the implanted electrode to a 16-channel headstage. Signals were streamed into a commercial acquisition system through a torqueless, feedback-controlled commutator (Tucker Davis Technology) and band-pass filtered between 100 and 5,000 Hz. Digital infrared videos of animal behavior from both side-and top-view cameras were simultaneously recorded at 640×480 pixel resolution at 25 frames per second (Streampix, Norpix). Video frame acquisition was triggered by a TTL pulse from the acquisition system to achieve synchronization between the video and the electrophysiological recording. Spikes were sorted manually using commercial software (OfflineSorter, Plexon) based on principal component analysis. Unit isolation was verified using autocorrelation histograms. To consider the recorded cell as a single unit, cells had to have a signal/noise ratio >2; spike shape had to be stable throughout the recording; and the percentage of spikes occurring with inter-spike intervals (ISIs) <3 ms (the typical refractory period for a neuron) in a continuous recording sequence had to be <0.1%. We checked for redundancies within days by examining the cross correlations of co-recorded neurons and checked for redundancies across days by comparing waveforms and temporal response profiles. After the first recording, the implanted electrode was slowly moved down in 40-µm increments. The placement of the electrode was examined histologically with the aid of DiI coated on the electrodes. Animals were excluded if electrodes were not confined to the PAG. Recordings of the VMHvl (Fig 1H-L) were performed previously using identical methods (Falkner et al. 2014; Lin et al. 2011) and reanalyzed here for direct comparison to PAG neurons.

### Electrophysiology Analysis

Spikes in single neurons were convolved with a 25 ms Gaussian for presentation (Fig 1D). Responsivity index for each behavior (Fig 1i-j) was computed as

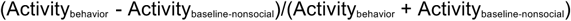

Where Activity_behavior_ is defined as the mean activity across all episodes of a particular behavior (e.g. attack) and Activity_baseline-nonsocial_ is defined as the mean activity across all episodes designated as non-social within the given social interaction. Within-neuron significance was determined using a paired t-test for each neuron (behavior vs. nonsocial) compared to a Bonferroni-corrected threshold for each tested population.

### PRV injections

The right masseter was exposed and PRV-152 (kind gift from Lynne Enquist), was injected at 5 separate locations, 250uL per injection along the A-P axis of the muscle. The skin was sutured closed and PRV was incubated for 96-112 hrs prior to sacrifice. In some cases, animals were allowed to freely interact with a male mouse for 10 min 1 hour prior to sacrifice at the 96 hr time point post-PRV injection. These animals were stained for c-Fos (primary antibody: goat anti-Fos, Santa Cruz, sc-52G, 1:300, secondary antibody: donkey anti-goat Dylight 549, Jackson Immuno, 705-505-147, 1:300).

### EMG implantation and recording

To perform simultaneous recordings of PAG neurons and jaw muscle activity, we implanted animals with chronic EMG electrodes in the right masseter superficial muscles of the jaw. Electrodes were constructed using a pair of 0.001 inch flexible multi-strand stainless steel wires (A-M Systems, No. 793200) with the insulation removed from a 0.5-mm segment of each wire such that pairs of electrodes recorded signals from separate but nearby areas of the same muscle. Electrode wires were threaded through the muscle during a surgical procedure and anchored with a knot on the outside of the muscle. EMG wires were then threaded under the skin to the base of the skull where they were attached to ground electrodes. EMG wire output was relayed through a preamplifier and commutator to the digitizer with a sampling rate of 3,000 Hz (Tucker Davis Technology). Signals were processed by taking the difference from the pair of electrodes, and this differential signal was low pass filtered at 300 Hz.

### EMG analysis

#### Mutual Information

Mutual information was computed between simultaneously recorded PAG activity and jaw EMG. EMG signals were rectified, low pass filtered at 20Hz, and downsampled to 1kHz. Spike trains were converted into a continuous instantaneous firing rate (IFR) with the same number of points as the downsampled EMG signals. For each pair of recorded PAG instantaneous firing rate and EMG signal, the continuous signals were discretized and MI was computed according to the definition(Shannon and Weaver 1964; Schilling, n.d.; Timme and Lapish 2018):

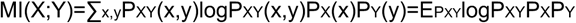

Where x and y represent the instantaneous firing rate and EMG signal respectively. For each signal pair, the MI was compared relative to the mean of ten iterations of a circularly permuted time shuffled control.

#### Spike Triggered EMG

STEMGs were computed on rectified EMG signals by averaging the EMG signal in an 800ms window around each PAG spike recorded during interactions with a male or with a female. STEMGs were computed separately for each 10s increment during male and female interactions, smoothed with a 5ms moving average, then normalized by the number of spikes. To correct for drift due to volleys of successive spikes, STEMGs were baseline corrected by subtracting a baseline 100ms boxcar filtered version of the STEMG. To determine whether STEMGs contained significant peaks, we set strict criteria: STEMGs had to have a minimum of 5 consecutive points that crossed above the 98% confidence interval within 60ms of the 0 (the spike onset).

## *In vitro* electrophysiological recordings

Vglut2:Cre x Ai6 mice were injected with 100 nL rAAV2.syn.flex.ChrimsonR.tdT into VMHvl. Three weeks after virus injection, acute horizontal brain slices of VMHvl and PAG (275 µm in thickness) were collected using standard methods (Fang et al. 2018). After being anesthetized by isoflurane inhalation, the mice were perfused by ice-cold choline based cutting solution containing (in mM) 25 NaHCO_3_,25 glucose, 1.25 NaH_2_PO_4_, 7 MgCl_2_, 2.5 KCl, 0.5 CaCl_2_, 110 choline chloride, 11.6 ascorbic acid, and 3.1 pyruvic acid. The slices were collected in the same cutting solution using a Leica VT1200s vibratome, incubated for 20 min in oxygenated artificial cerebrospinal fluid (ACSF) solution (in mM: 125 NaCl, 2.5 KCl, 1.25 NaH_2_PO_4_, 25 NaHCO_3_, 1 MgCl_2_, 2 CaCl_2_ and 11 glucose) (osmolality, 295 mmol/kg) at 32– 34°C and then maintained at room temperature until use. Standard whole cell recordings were performed with MultiClamp 700B amplifier (Molecular Devices) and Clampex 11.0 software (Axon Instruments). Membrane currents were low-pass filtered at 2 kHz and digitized at 10 kHz with Digidata 1550B (Axon Instruments). Electrode resistances were 2–4 MΩ, and most neurons had series resistance from 4 to 15 MΩ. Glutamatergic (green fluorescent) or GABAergic (non-fluorescent) cells as well as Chrimson-tdTomato expressed VMHvl cells were identified with an Olympus 40 x water-immersion objective with GFP and TXRED filters. The slices were superfused with ACSF warmed to 32– 34°C and bubbled with 95% O_2_ and 5% CO_2_. The intracellular solutions for voltage clamp recording contained (in mM) 135 CsMeSO_3_, 10 HEPES, 1 EGTA, 3.3 QX-314 (Cl– salt), 4 Mg-ATP, 0.3 Na-GTP, and 8 Na_2_-Phosphocreatine (osmolality, 295 mmol/kg; pH 7.3 adjusted with CsOH), and for current clamp recording contained (in mM) 130 K MeSO_3_, 5 KCl, 0.5 EGTA, 20 HEPES, 1.8 MgCl_2_, 0.1 CaCl_2_, 4 Na_2_-ATP, and 0.2 Na-GTP (osmolality, 295 mmol/kg; pH 7.3 adjusted with KOH). To activate Chrimson-expressing VMHvl glutamatergic neurons and Chrimson-expression axons in PAG, brief pulses of full field illumination (20 ms for VMHvl during current clamp recording and 1 ms duration for PAG during voltage clamp recording) were delivered onto the recorded neuron with 605 nm LED light (pE-300white; CoolLED) at 35 s intervals. Voltage clamp recording was conducted for PAG neurons, and the membrane voltage was held at −70 mV for EPSC recording, and at 0 mV for IPSC recording. Current clamp recording was conducted in VMHvl glutamatergic neurons expressing Chrimson-tdTomato, where the neurons were maintained at resting potential and spiking activity was detected with or without red light pulses (20 ms, 20 Hz for 500 ms).

## Fiber photometry recordings

A rig for performing simultaneous fiber photometry recordings from 2 locations was constructed following basic specifications previously described with a few modifications. For simultaneous VMH and VMHvl-PAG recordings, we injected vGlut2-ires-cre males with 100-160nl of AAV1.CAG.Flex.GCaMP6f.WPRE.SV40 (Lot CS0956, CS0845, CS0224WL, Upenn, final titer:: 9.3 × 10^12 PFU/ml) ipsilaterally into the VMHvl, and 240nl of HSV-Ef1a-LS1L-GCaMP6f (MIT vector core,Lot RN506, final titer: 1.0 × 10^9 PFU/ml) contralaterally in the PAG. For simultaneous recordings of the VMHvl and lPAG, we injected 80-120nl of AAV2/1 CAG::Flex-GCaMP6f-WPRE-SV40 ipsilaterally into the VMHvl, and 160-240nl of AAV2/1 CAG::Flex-GCaMP6f-WPRE-SV40 in the lPAG. VIruses were injected using the following coordinates: VMHvl (−1.82A/P, 0.72M/L, −5.8D/V), lPAG(−3.64A/P, 0.5M/L-2.4D/V).

A 400-μm optic fiber (Thorlabs, BFH48-400) housed in a metal ferrule (Thorlabs, SFLC440-10) was implanted 0.4 mm above each injection site, except for the HSV-Ef1a-LS1L-GCaMP6f injection, where the fiber was placed over the VMHvl. After three weeks of viral incubation and before recording, a matching optic fiber was connected to the each implanted fiber using a ferrule sleeve. A 400-Hz sinusoidal blue LED light (30-50 μW) (LED light: M470F1; LED driver: LEDD1B; both from Thorlabs) was bandpass filtered (passing band: 472 ± 15 nm, Semrock, FF02-472/30-25) and delivered to the brain to excite GCaMP6. The emission light then traveled through the same optic fiber, was bandpass filtered (passing band: 534 ± 25 nm, Semrock, FF01-535/50), detected by a femtowatt silicon photoreceiver (Newport, 2151) and recorded using a real-time processor (RZ5, TDT). The envelope of the 400-Hz signals that reflects the intensity of the GCaMP6 signals were extracted in real-time using a custom TDT program. Baseline adjusted fluorescence signals were regressed using a 30s spline approximation.

## Histology and imaging

Animals were deeply anaesthetized using 0.5 ml of a ketamine-xylazine cocktail (10 mg/ml ketamine and 5 mg/ml xylazine) and transcardially perfused with phosphate buffered saline (PBS) followed by cold 4% paraformaldehyde in PBS. Brains were immersed overnight in a 20% sucrose solution, embedded with cutting medium (Tissue-Tek) and sectioned using a cryostat (Leica). Standard immunohistochemistry procedures were followed to stain 30-μm coronal brain sections for all mice. DAPI (1:20,000, Life Technologies, catalog number D21490, widely validated) was used to assess electrode track for physiology and fiber track for photometry. We acquired 2.5× or 5× fluorescent images to determine cannula or electrode placements. We used 10× fluorescent images to count c-Fos+ and PRV+. Cell counting was done manually using ImageJ on 30-μm sections separated by 60 μm that had observed GFP label in the lPAG.

## Time-varying linear regression model

We modeled the population response of the PAG during male and female interactions for each animal by fitting the PAG response during each interaction including 10s prior to the introduction and 10s following the removal of the animals with a series of autoregressive models using time-varying lengths of simultaneously recorded VMHvl signal as the variable regressors using the form:

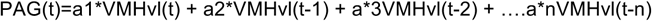

where t represents the current time bin and the other regressors represent variable amounts of elapsed time. PAG and VMHvl signals were binned in either 50ms, 100ms, 250ms, 500ms, and 1s bins and models were fit independently for each of these bin sizes. For each VMHvl and PAG signal, data was halved and the first half was used to fit and the second half was used to cross validate, using least squares. Model order was set to maximum value of 10s of elapsed time, set independently for each bin size. For each family of models fit to the data, the best order model was determined using AIC (Akaike Information Criteria) on the fit to the cross validated data. A timeshifted null model was fit by circularly permuting the input data (VMHvl) signal and fitting the unpermuted PAG signal, and fit and model order were re-fit for the timeshifted data. Model order for the selected model was converted to elapsed time for each binsize. Fits were performed separately for male and female interactions. Additionally, we tested the alternative hypothesis that PAG signals influence VMHvl signal by fitting the “reverse” model of the form:

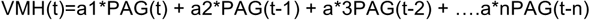

A similar model family was fit for the reverse model, and the model order (and elapsed time) was determined for each model independently for each bin size.

## Statistical Analysis

Parametric tests, including Student’s *t*-test, paired *t-*test, and two-sample *t* test were used if distributions passed Kolmogorov–Smirnov tests for normality. For within-neuron tests of firing rate significance, a non-parametric Wilcoxon signed rank test was used since spike rates were often low and not normally distributed. Repeated tests of significance were corrected with a strict Bonferroni correction. For all statistical tests, significance was measured against an alpha value of 0.05 unless otherwise stated. All error bars show s.e.m. No statistical methods were used to predetermine sample sizes, but our sample sizes are similar to those reported in previous publications. Data collection and analysis were not performed blind to the conditions of the experiments.

Statistical analyses used in each figure are listed below.

Figure 1. (B) Single factor, repeated measures ANOVA. (D) Single neuron activity average using 25ms Gaussian smoothing (E-G) Top panels are the z-scored responses of individual neurons aligned to each behavior, and histograms represent the number of neurons whose peak response lies at that bin relative to behavior onset. Dotted line represents chance level (number of bins/number of neurons). (I-J) Wilcoxon signed rank test for comparison of population response. Percentages of neurons in pie charts computed using within-neuron significance test across one or both behaviors, with Bonferroni correction. (K) Kolmogorov-Smirnov test (L) Unpaired ttest across all bins using Bonferroni corrected threshold across all bins.

Figure 2. (C) Paired t-test, (f) Kolmogorov-Smirnov test between cumulative distributions. (G) Significant EMGsm+ neurons see methods.

Figure 3. (F) Fisher’s test, (H) paired t-test, (M-Q) Error bars show +/−SEM, comparisons of means with paired t-test. (U) Error bars show +/−SEM (V) Wilcoxon signed rank test of male, female, and baseline activity. (W) Paired t-test between forward and null model at each bin for fit, Paired t-test between male and female interactions for model time.

Supplementary Figure 1. (A-F) Paired t-test.

Supplementary Figure 2. (C) Paired t-test.

Supplementary Figure 3. (A-C) Wilcoxon signed rank test.

Supplementary Figure 4. (A-B,D-E) Activity shown is z-scored across the whole interaction trace. (C,F) Activity for each individual behavior is baseline subtracted using a 1s bin 5s prior to interaction. Individual behaviors compared using student’s ttest, and comparison between behaviors using an unpaired t-test.

Supplementary Figure 5. (A-B) Paired t-test between forward and reverse model at each bin.

Supplementary Movie 1. Pharmacological inactivation of the PAG using muscimol results in aggression-specific deficits. Example intermale aggression following saline infusion, compared with behavior following muscimol inactivation. During inactivation, intermale aggression is decreased while investigatory behaviors and other social behaviors are unchanged.

Supplementary Movie 2. Example of single lPAG neuron recorded simultaneously with EMG in the superficial masseter muscle of the jaw with corresponding behavior. lPAG neuron (bottom trace) shows time-locked spiking with EMG peaks (top trace).

---

**Supplementary Figure 1.**
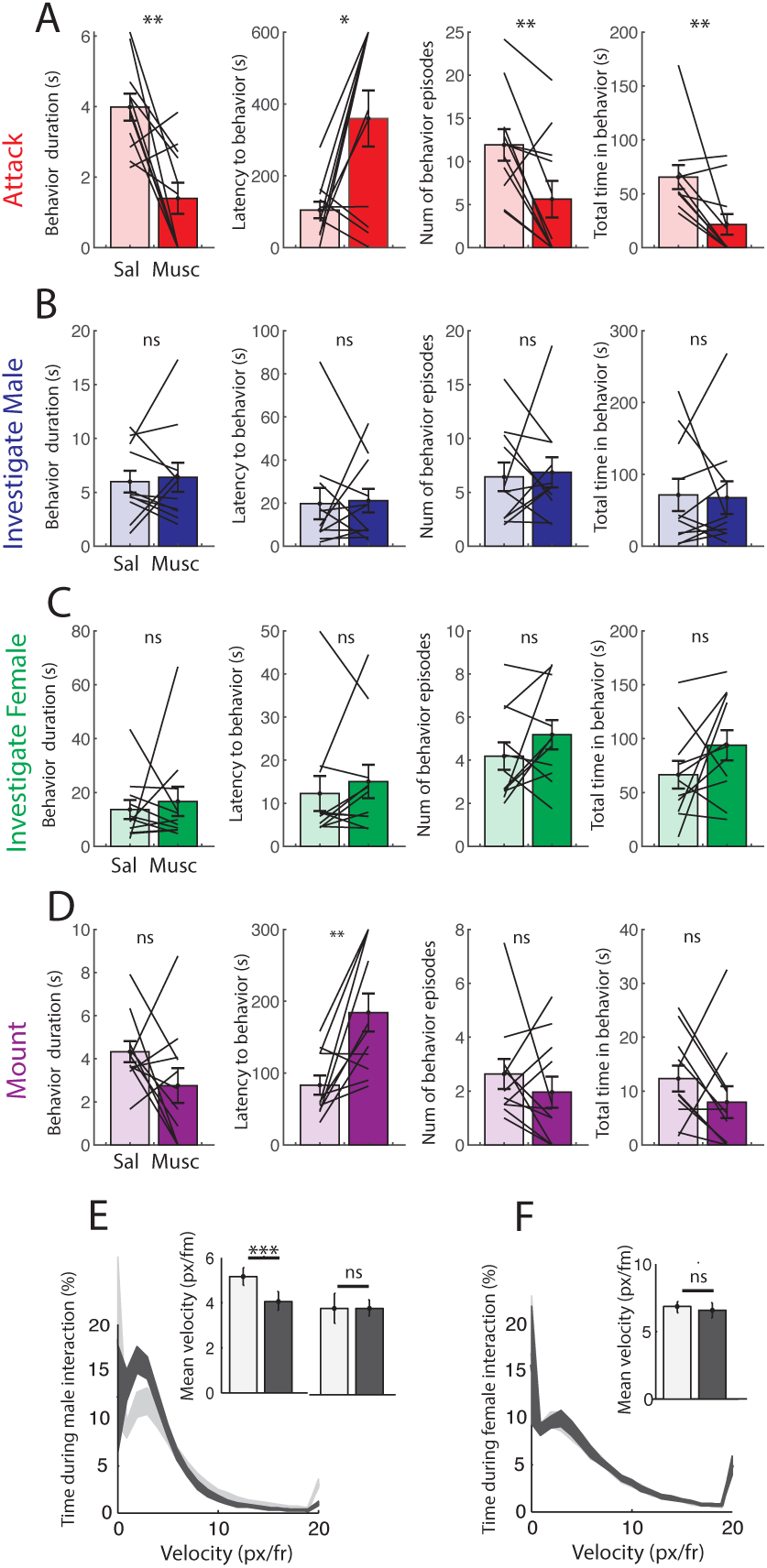
Alternative measures of behavior, including behavior duration, latency to first behavior, number of behavior episodes, and total time spent engaging the behavior confirm reduction of aggression. Behaviors shown are attack (A, **p=0.00284, *p=0.0121, **p=0.0090,**p=.00689), investigation of male (B, p=0.7344, p=0.8617, p=0.8091, p=0.8719), investigation of females (C, p=0.6784, p=0.4077, p=0.2029, p=0.1322), and mounting behavior (D, p=0.1490, **p=0.0036, p=0.3830, p=0.2329). Mean saline shown in light bars, muscimol shown in dark bars. Reversible inactivation during male interaction reduces mean resident velocity (E), but effects are eliminated when episodes of attack are removed from analysis (E, right inset). Inactivation during female interaction has no effect on resident velocity (F). (N=11 animals for all comparisons, *p<0.05, **p<0.01, ***p<0.001, ns=nonsignificant, paired ttest).

**Supplementary Figure 2.**
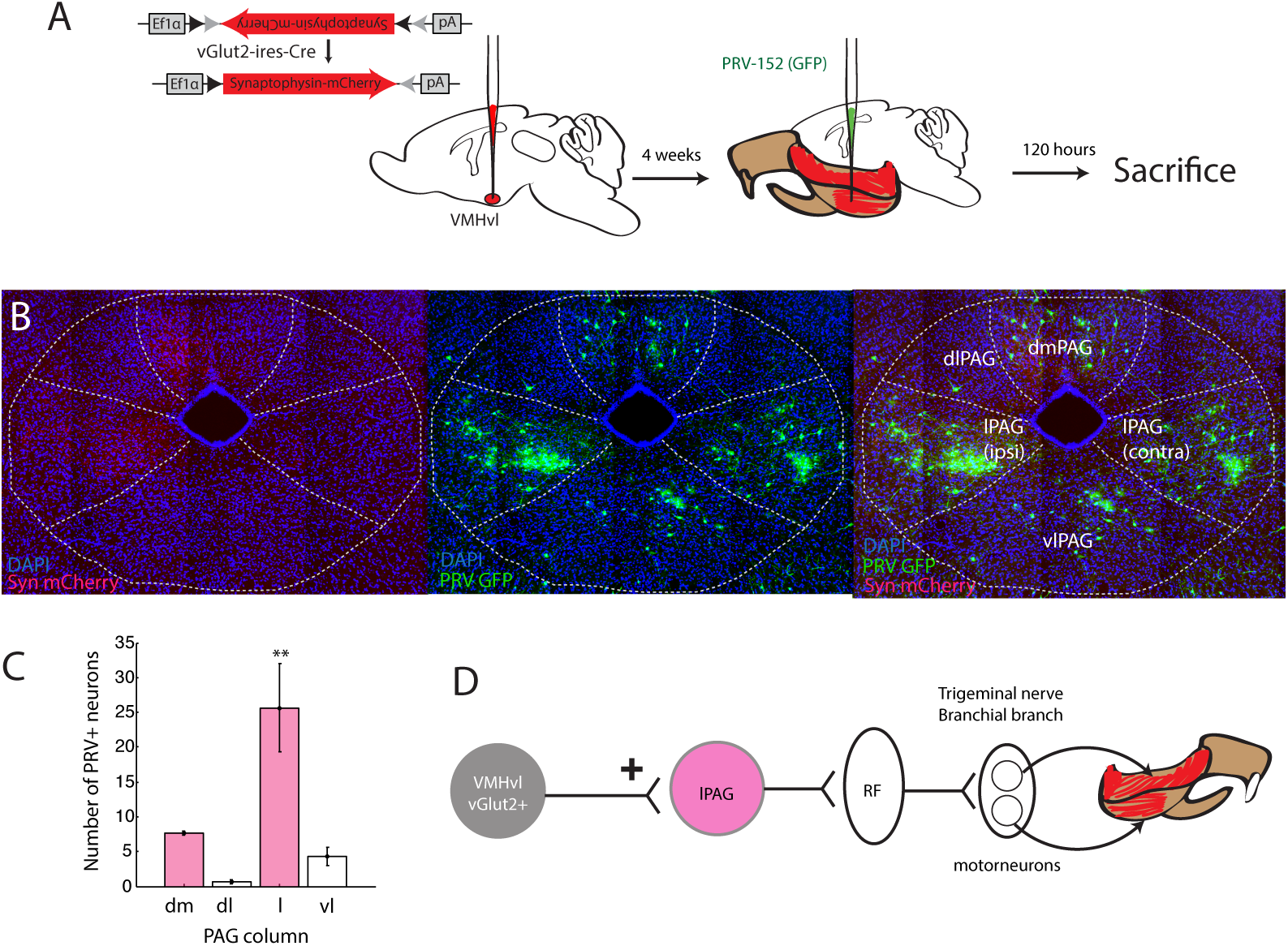
PRV-labeled jaw-projecting neurons are maximally within VMHvl-projection-defined column boundaries. (A) Experimental procedure for determining overlap between VMHvl projection fields and PRV label. (B) Example histology showing VMHvl synaptophysin-containing terminals (left), PRV-GFP labeled jaw-projecting neurons (center), and overlap (right). (C) Number of GFP labeled neuron observed within each PAG column. N=4 animals, p=0.0028, one-way ANOVA (D) Putative circuit from excitatory vGlut2+ neurons to jaw.

**Supplementary Figure 3.**
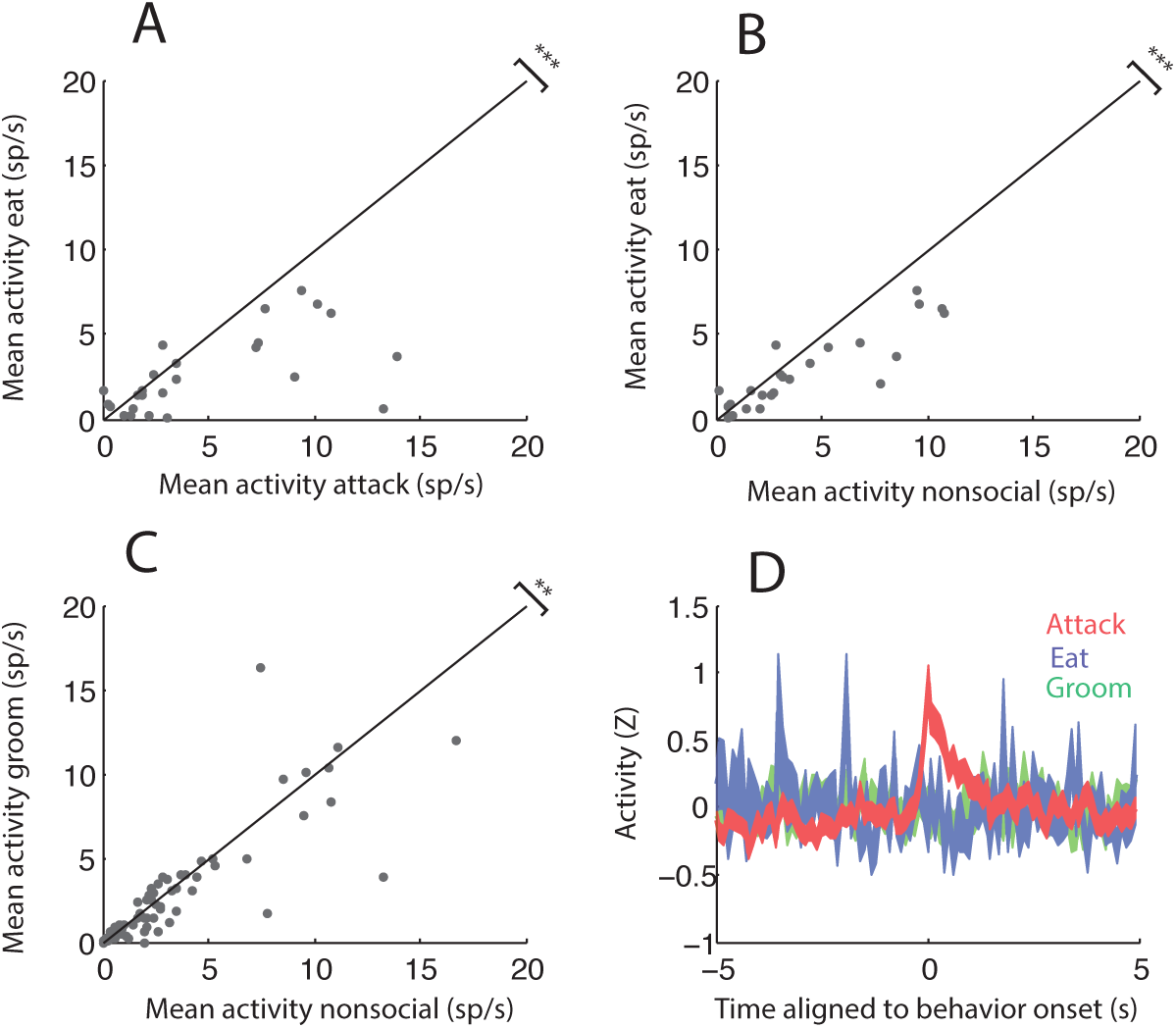
Other jaw-related behaviors decrease lPAG activity. (A-B) Activity during eating is significantly decreased relative to attack (A, N=26, p=0.0007), and nonsocial epochs (B, N=26, p=0.0006). Gray dots represent single neurons tested in both behaviors. (C) Activity during grooming is decreased relative to nonsocial epochs (N=87, p=0.004). (D) Neurons with significant STEMG activity do not show activity increases aligned eating or grooming onsets, but do show activity aligned to attack. All pairwise comparisons done with Wilcoxon signed rank test.

**Supplementary Figure 4.**
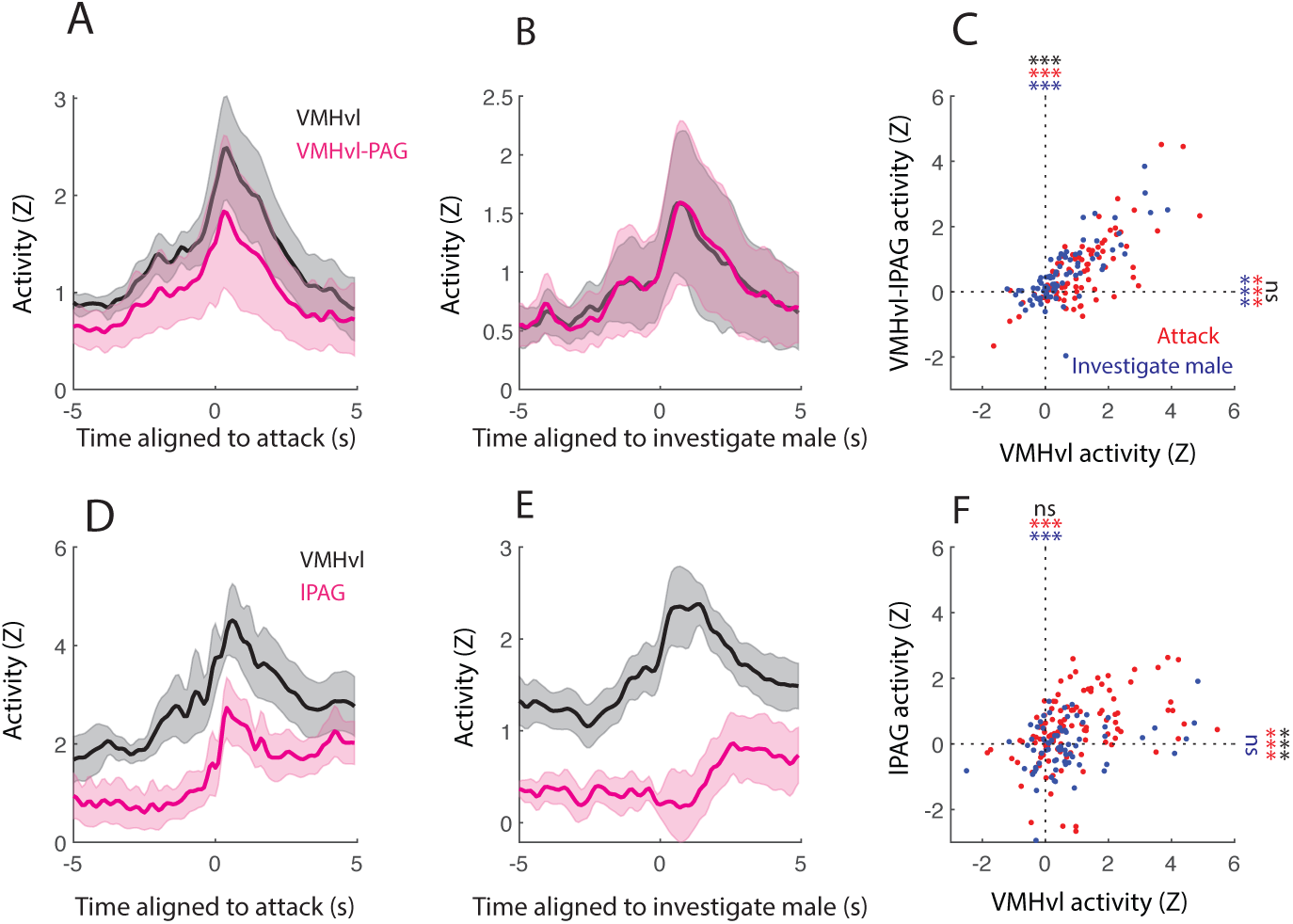
Fiber photometry responses to individual male behaviors during simultaneous recordings. (A-B) PETHs comparing simultaneously recorded mean VMHvl (black) and VMHvl-lPAG (magenta) aligned to attack (A) and investigate male (B) N=6 animals. (C) Comparison of individual behavior episodes for attack (red) and investigate male (blue). Both attack and investigate male acutely increase VMHvl and VMHvl-lPAG responses (***p=9.2157*10^-13, ***p=1.3144*10^-6, N=84 attack episodes in 6 animals; ***p=1.8445*10^-7, ***p=2.6243*10^-7, N=85 investigate male episodes in 6 animals, one sample ttest; Comparison between behaviors, VMHvl: ***p=5.157*10^-4, VMHvl-lPAG: p=0.4201). (D-E) PETHs comparing simultaneously recorded mean VMHvl (black) and lPAG (magenta) aligned to attack (D) and investigate male (E) N=7 animals. (F) Comparison of individual behavior episodes for attack (red) and investigate male (blue). Attack acutely increases VMHvl and lPAG (***p=2.9198*10^-11, ***p=4.0921*10^-5, N=84 attack episodes in 6 animals), but investigate only increases VMHvl (***p=2.1263*10^-5, p=0.5164, N=85 investigate male episodes in 7 animals, one sample ttest). Comparison between behaviors, VMHvl: ***p=5.157*10^-4, VMHvl-lPAG: p=0.4201). lPAG responses are significantly different between attack and investigate, while VMHvl responses are not (VMHvl: p=0.1467, lPAG: ***p=5.7677*10^-4, N=84 attack episodes, N=85 investigate male episodes in N=7 animals, unpaired ttest).

**Supplementary Figure 5.**
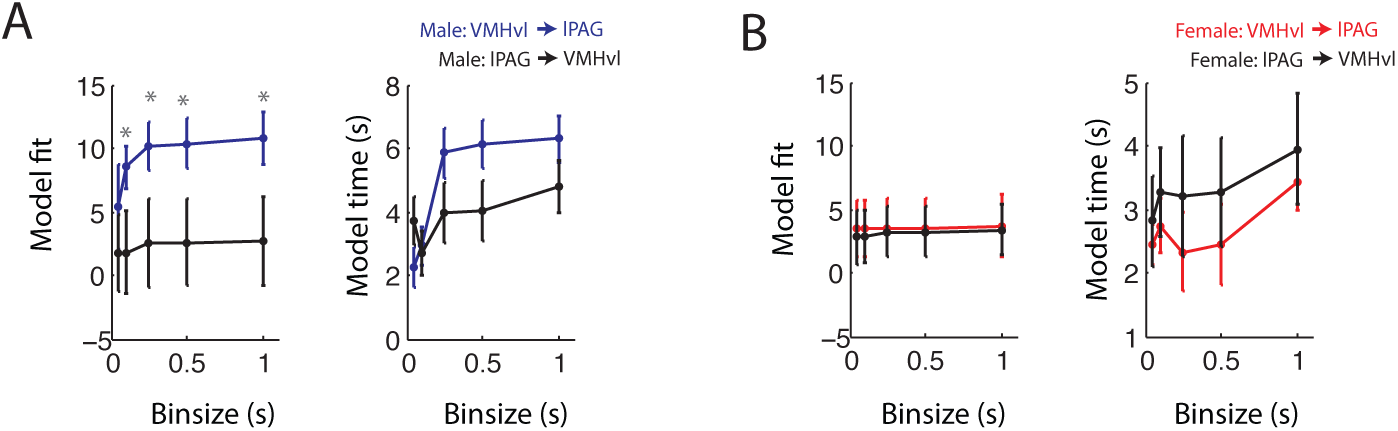
Comparison of forward (VMHvl → lPAG) and reverse (lPAG ? VMHvl) regressive models. Model fit percentages and model time for fits during male (A) and female (B) interactions. Black traces (A-B) show reverse models. During male interactions (blue), fits are significantly increased in forward model relative to reverse model (*p<0.05, paired t-test for each time bin).

## Author contributions

ALF designed and carried out *in vivo* physiology and photometry experiments, performed all analysis and modeling for these experiments, and wrote the manuscript. DW conducted slice physiology experiments, analyzed the data and co-wrote the manuscript. AS assisted with PRV tracing experiments, LWW and IZC performed pharmacological inactivation experiments. JEF constructed microdrives associated with in vivo physiology experiments. DL conceived the project, suggested experiments, analyzed data and edited the manuscript.

